# Population genetic structure of the maize weevil, *Sitophilus zeamais*, in southern Mexico

**DOI:** 10.1101/2022.02.11.480163

**Authors:** Jennifer Baltzegar, Michael S. Jones, Martha Willcox, Janine M. Ramsey, Fred Gould

**Affiliations:** Genetic Engineering and Society Center, North Carolina State University, Raleigh, NC, USA; Graduate Program in Genetics, North Carolina State University, Raleigh, NC, USA; International Maize and Wheat Improvement Center (CIMMYT), Texcoco, México; Instituto Nacional de Salud Pública/CRISP, Tapachula, Chiapas, México; Department of Entomology and Plant Pathology, North Carolina State University, Raleigh, NC, USA

## Abstract

The maize weevil, *Sitophilus zeamais*, is a ubiquitous pest of maize and other cereal crops worldwide and remains a threat to food security in subsistence communities. Few population genetic studies have been conducted on the maize weevil, but those that exist have shown that there is very little genetic differentiation between geographically dispersed populations and that it is likely the species has experienced a recent range expansion within the last few hundred years. While the previous studies found little genetic structure, they relied primarily on mitochondrial and nuclear microsatellite markers for their analyses. It is possible that more fine-scaled population genetic structure exists due to local population dynamics, the biological limits of natural species dispersal, and the isolated nature of subsistence farming communities. In contrast to previous studies, here, we utilized genome-wide single nucleotide polymorphism data to evaluate the genetic population structure of the maize weevil from the southern and coastal Mexican states of Oaxaca and Chiapas. Our study is the first to find significant genetic population structure in the maize weevil. Here, we show that although there continues to be gene flow between populations of maize weevil, that fine-scale genetic structure exists. It is possible that this structure is shaped by the movement and trade of maize by humans in the region, by geographic barriers to gene flow, or a combination of factors.

## Introduction

The maize weevil, *Sitophilus zeamais*, is a ubiquitous pest of maize and other cereal crops worldwide. Although there are effective control solutions, these are not always available to subsistence farmers. Therefore, maize weevils remain a threat to food security in these communities. Recent research to control the maize weevil has focused on developing maize varieties resistant to the insect, improving storage conditions, and identifying chemical compounds toxic to the species [e.g., 1-3]. Maize varieties that are resistant to damage tend to have higher levels of phenolic compounds, which increases the hardness of grain and provides mechanical resistance to the maize from the maize weevil [4]. However, subsistence farmers, especially in Mexico, often prefer to grow their own landraces, which are maize varieties adapted for the specific climate, taste, and texture preferences in the region and may not have been specifically selected to protect against storage pests [5]. Hermetic storage solutions can be very effective, but they can also be prohibitively expensive for subsistence farmers. Recently, advances in more affordable hermetic storage bags have been made and programs exist to give access to silos and these improved storage solutions. However, silos and other hermetic storage containers are still not available to all who need them. Another major area of research aimed to control the maize weevil is in the discovery of chemical agents, especially plant-derived essential oils, that may help to reduce population sizes in stored grain. It remains unclear if these types of chemicals can be produced in quantities that are large enough and affordable for widespread adoption [6]. In addition, insecticide resistance has been reported in this species and resistance to any new chemical control agents would be expected to develop with sustained use, although some posit that resistance to essential oils would develop more slowly than resistance to synthetic pesticides owing to the complex chemical makeup of essential oils [6].

Few population genetic studies have been conducted on the maize weevil, but those that exist have shown that there is very little genetic differentiation between geographically dispersed populations. Eight nuclear microsatellite markers were previously used to assess 15 populations from 9 countries over 4 continents [7]. Their conclusion was that very little genetic structure exists globally. A second study used mitochondrial markers to identify population structure among 9 countries in West and Central Africa [8]. Again, their conclusion was that very little phylogeographic genetic structure exists in this species. Together, these results are evidence for the recent worldwide expansion of the maize weevil.

While a recent expansion of the maize weevil has likely occurred within the last few hundred years, it is also probable that more fine-scaled population genetic structure exists due to local population dynamics, the biological limits of natural species dispersal, and the isolated nature of subsistence farming communities. The typical flight range for maize weevil was roughly estimated to be one quarter mile [9]. In addition, subsistence farmers are likely not trading grain at an international scale, thus genetic differences in maize weevil may be found between these types of communities.

Mitochondrial and nuclear microsatellite markers are classic methods to assess the population genetic structure in species of interest [10]. They are especially good at identifying population structure that has existed for a significant period of time. However, the use of so few markers limits the statistical power that these studies have to identify recently developed genetic structure. In these cases, the finding of no structure is not proof for the absence of structure, rather it is likely that these studies are underpowered [10]. It is prudent to utilize a more powerful technique to identify possible genetic structure. Single nucleotide polymorphisms (SNPs) are a marker-type that have higher statistical power to answer these types of questions and are used regularly to elucidate population structure. One individual bi-allelic SNP is less powerful than an individual microsatellite because of the fewer number of alleles sampled. But, when hundreds or thousands of SNPs are sampled at once, the combined statistical power is higher than the few dozen microsatellite loci that may be available for a species [11]. Maize weevil has only 8 polymorphic and amplifiable microsatellite loci identified [7].

Historically, it was challenging to generate a large number of informative SNPs for a non-model organism. However, reduced genome representation sequencing methods allow researchers to sample genome-wide SNPs without the need for *a priori* knowledge of the genome. Reduced representation methods have given researchers a greater ability to characterize the genetic population structure of important, but non-model organisms. Here we use double-digest restriction-site associated sequencing (ddRadSeq) to identify informative SNPs and assess the genetic population structure of the maize weevil collected from subsistence communities in southern Mexico.

Mexico has been chosen as the study location as it is near the center of origin for maize and the maize weevil is the most common pest present in stored maize in Oaxaca and Chiapas. The region continues to have strong subsistence farming communities and each community grows landraces of maize specific to their region and preferences. Human mediated transport via the grain trade is likely the most common way that maize weevils migrate. The specific amount of trade between subsistence households in Mexico is not well characterized. Natural migration via flight is also a possible method of migration. Oaxaca and Chiapas are separated by the Isthmus of Tehuantepec. The isthmus is the shortest land distance between the Pacific Ocean and the Gulf of Mexico and is a great source for wind energy [12]. Given that maize weevils typically fly short distances (∼0.25 miles) and are not particularly strong fliers, the isthmus may provide a barrier to gene flow [9]. In addition, both states exhibit a wide range of elevations and climates, providing further opportunity for local adaptation to occur and population structure to be identified. It is imperative to better understand the basic population dynamics underlying this species so that traditional control methods can better be applied. Furthermore, the information in this study will be vital for informing the socio-ecological context in which maize weevil exists, allowing researchers who might develop emerging technologies for maize weevil control to identify relevant stakeholders from which to seek approval [13].

## Materials and Methods

### Field collections

From 2 August to 18 August 2016, individual households in the Mexican states of Oaxaca and Chiapas were visited. Rural communities and small to medium sized farms within these communities were chosen based on existing relationships with community members that had been previously developed. At each residence, we recorded latitude, longitude, and elevation using a Garmin GPS unit (Garmin International, Inc., Olathe, KS, USA). Individual maize weevils from stored maize were collected, if the pest was present. These samples were stored in 100% ethanol at ambient temperature during the duration of field collecting. Upon returning to the United States, samples were stored at -20°C in the laboratory until processed.

### Genomic DNA isolations

Genomic DNA was isolated from the head and thorax of individual weevils following the DNeasy Qiagen Kit (Qiagen Inc., Germantown, MD, USA) manufacturers suggestions. Samples were incubated overnight in the lysis buffer solution at 55°C and on day two of the isolation a RNaseA (4mg/mL) treatment was performed before the recommended wash steps. Finally, samples were eluted into 300 µl of 70°C dH_2_O and stored at -20°C for short term storage or - 80°C for long term storage.

### ddRadSeq library preparation and sequencing

Individual weevils were prepared for sequencing following the ddRadSeq protocol first described by Peterson *et al*. [14] and adapted to our laboratory as described by Fritz *et al*. [15]. Two separate libraries were prepared and sequenced on two different lanes of an Illumina HiSeq 2500 125 bp SE. The first library included 96 individual weevils, 24 each from the 4 most distant communities sampled: Santa Maria Yavesia, Santa Rosa de Lima, Huixtla, and Nuevo Leon. This sampling regime included the communities with the highest and lowest elevation from each state. This particular library was used to test the hypothesis that genetic differentiation could be found by genotyping SNPs. A second library, sequenced in the same manner as the first, was prepared after confirmation that genetic differentiation could be identified. This library included 12 individual weevils from each of the remaining 8 cities sampled, Santa Ana Zegache, Santos Reyes Nopala, San Isidro Campechero, Santiago Yaitepec, Pochutla, Golondrinas, Montecristo, and Villa Corzo.

### Bioinformatic and statistical analyses

Data produced by each lane was first checked with FastQC, then reads were trimmed with trimmomatic to remove adapter and Illumina indices [16-17]. The trimming was verified by FastQC and a multiQC report was generated to compare results from the different indices [16,18]. Data from both lanes were analyzed together in Stacks v. 1.48 following the *de novo* procedure described by Rochette and Catchen [19] [20]. The procedure is outlined here to specify certain parameter choices. Samples were first cleaned, demultiplexed, and truncated to 115 bp using the program *process_radtags*. The per sample coverage was checked. Then, *ustacks* was run on all samples (-M = 5, -m = 3). The 4 highest coverage individuals from each of the 12 populations were chosen to create the catalog, with the exception of sample zag.c1, which had much higher coverage than other samples. Using the catalog popmap created in the previous step, *cstacks* was implemented (-n = 5). Then, *sstacks* was run on all samples. To improve calls, *rxstacks* was used with the flags --prune_haplo and --conf_lim = 0.10. Then, *cstacks* and sstacks were run again using the improved catalog. Finally, genotype information for each individual was exported from Stacks in genepop format from the *populations* program. Using the *adegenet* package in R, these data were stringently filtered [21-22]. Individuals with greater than 80% missing genotype calls were removed. Then, loci with greater than 35% of missing data were removed. Principal components were then computed and plotted with the same program. F statistics were calculated in *hierfstat* package in R [23].

## Results

### Field collections

We sampled 7 communities in Oaxaca, Mexico and 5 communities in Chiapas, Mexico (Fig 1). In each city, we collected maize weevils from 3 to 7 individual residences and recorded environmental and biological parameters (Table 1). The sites in Oaxaca ranged in elevation from 23.5 m to 2,005 m and those in Chiapas ranged from 8.0 m to 1,142.5 m. When weevils were present, approximately 200 individuals were collected per residence. Weevils were collected from a total of 61 households.

**Table 1.** Summary table of communities visited in Oaxaca and Chiapas. Table includes the label for each community, which corresponds to figure labels, number of households visited, mean latitude and longitude for each community, and mean elevation (m) of households visited in each community.

| State | Community | Label | Homes | Elevation | Latitude | Longitude |
| --- | --- | --- | --- | --- | --- | --- |
| Oaxaca | Santa María Yavesía | YAV | 6 | 1988.25 | 17.234986 | -96.436065 |
|  | Santa Ana Zegache | ZEG | 4 | 1482.94 | 16.837518 | -96.724833 |
|  | Santos Reyes Nopala | NOP | 7 | 466.57 | 16.078580 | -97.154197 |
|  | San Isidro Campechero | ISD | 7 | 105.53 | 15.980961 | -97.302886 |
|  | Santiago Yaitepec | YAI | 5 | 1801.40 | 16.226998 | -97.268016 |
|  | Santa Rosa de Lima | SRL | 3 | 24.27 | 16.039082 | -97.491641 |
|  | San Pedro Pochutla | POC | 6 | 145.58 | 15.760327 | -96.499309 |
| Chiapas | Huixtla | HUX | 5 | 13.72 | 15.071078 | -92.543628 |
|  | Golondrinas | GOL | 4 | 745.00 | 15.419210 | -92.651981 |
|  | Montecristo | MTC | 4 | 329.31 | 16.937728 | -93.307673 |
|  | Villa Corzo | VLC | 5 | 585.65 | 16.181779 | -93.269561 |
|  | Nuevo León | NVL | 5 | 1106.10 | 16.486764 | -92.572740 |

**Fig 1.**
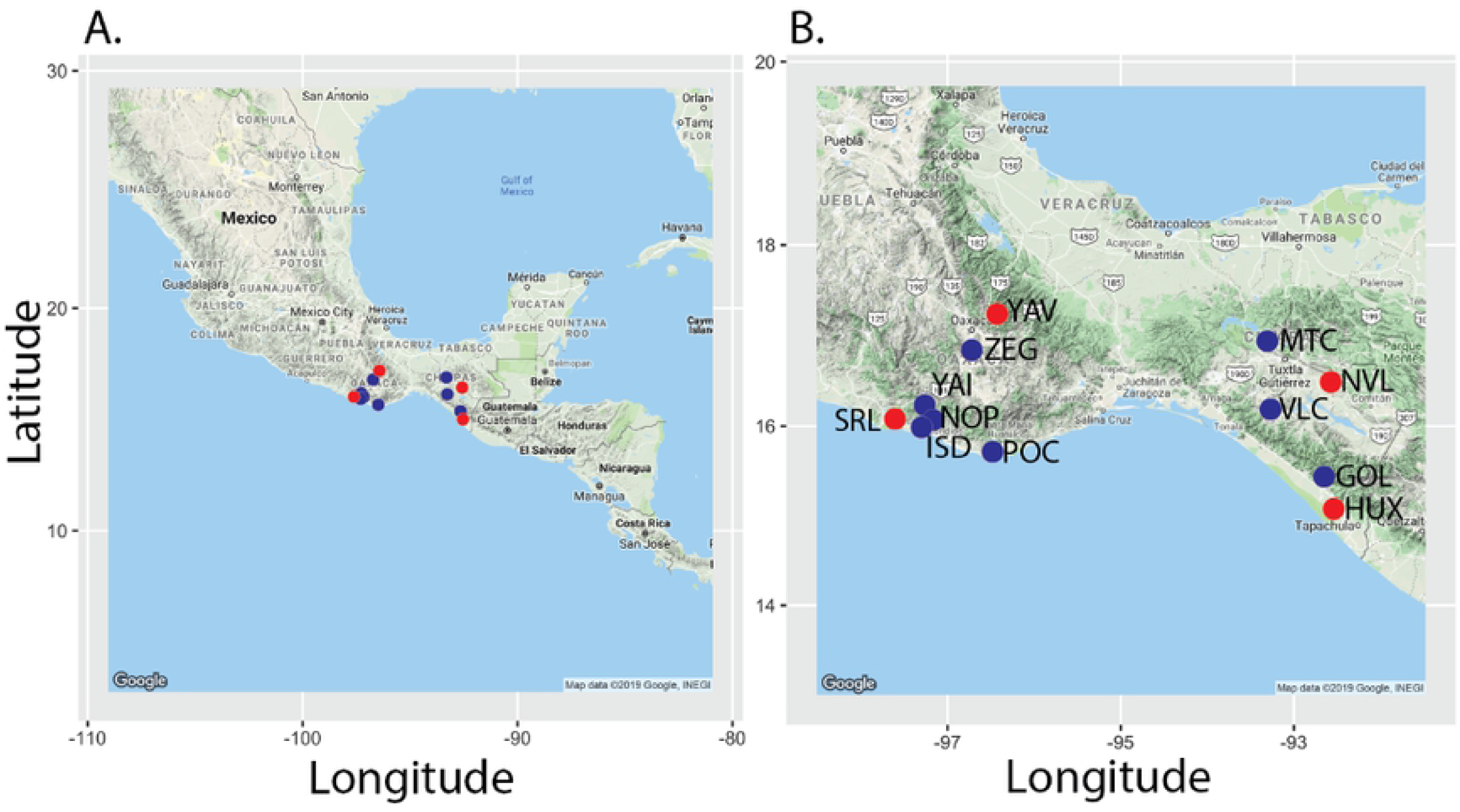
Map of sampling locations. Communities marked with red dots were sequenced in Lane 1, samples marked with blue dots were sequenced in Lane 2. Full community names are listed in Table 1. A) A map of Mexico showing relative location of sampled communities located in the southern and coastal region of Mexico. B) A zoomed in view centered on Oaxaca and Chiapas and the location of the communities where maize weevils were collected.

### Genomic DNA isolations

Twenty-four individual weevils were isolated from 4 communities for the proof-of-concept sequencing run. Twelve individuals each from the 8 remaining communities were isolated for the second sequencing run.

### ddRadSeq

The first sequencing run resulted in 111,204,487 reads after trimming and filtering with *process_radtags*, with an average 1,158,380 reads per individual and a range of 288,190 – 3,079,538 (Table 2). The second sequencing library resulted in 186,219,498 trimmed and processed reads, with a range of 15,068 – 4,730,073 and an average 1,939,786 reads per individual (Table 2). Although the initial quality check appeared similar in each sequencing run, there were clear sequencing lane differences upon inspection of the PCA plot produced after running *populations* with both lanes together in Stacks (Fig 2) therefore, stringent filtering criteria were applied. There were four populations, Santos Reyes Nopala (NOP), Golondrinas (GOL), Villa Corzo (VLC), and Nuevo Leon (NVL), where every individual had greater than 80% missing genotypes in the *populations* output, these were removed from further analysis. The populations with missing data were from both sequencing lanes. The mean number of reads for these populations following the *process_radtags* procedure was 1,680,432, while the mean number of reads for all populations was 1,549,083. Therefore, the reason for the high number of missing genotypes is likely due to specific parameter selection downstream of the *process_radtags* program. Following individual removal, a missing data threshold for individual SNPs was set at 35%, this resulted in the removal of 21,365 SNPs from the analysis. The final number of individuals included in the analysis was 132. The final number of SNPs included was 17,966.

**Table 2.** Sequencing results for two lanes of Illumina HiSeq 125 bp SE data. 192 individual weevils from 12 populations were prepared with the ddRadSeq protocol as described by Peterson *et al*. [14] and processed in Stacks v. 1.48. Table includes the total read count per lane, the read count for the individuals with the lowest and highest coverage per lane, and the mean number of reads per individual per lane.

| Lane | Total Reads | Low Read Count | High Read Count | Mean Reads Per individual |
| --- | --- | --- | --- | --- |
| 1 | 111,204,487 | 288,190 | 3,079,538 | 1,158,380 |
| 2 | 186,219,498 | 15,068 | 4,730,073 | 1,939,786 |

**Fig 2.**
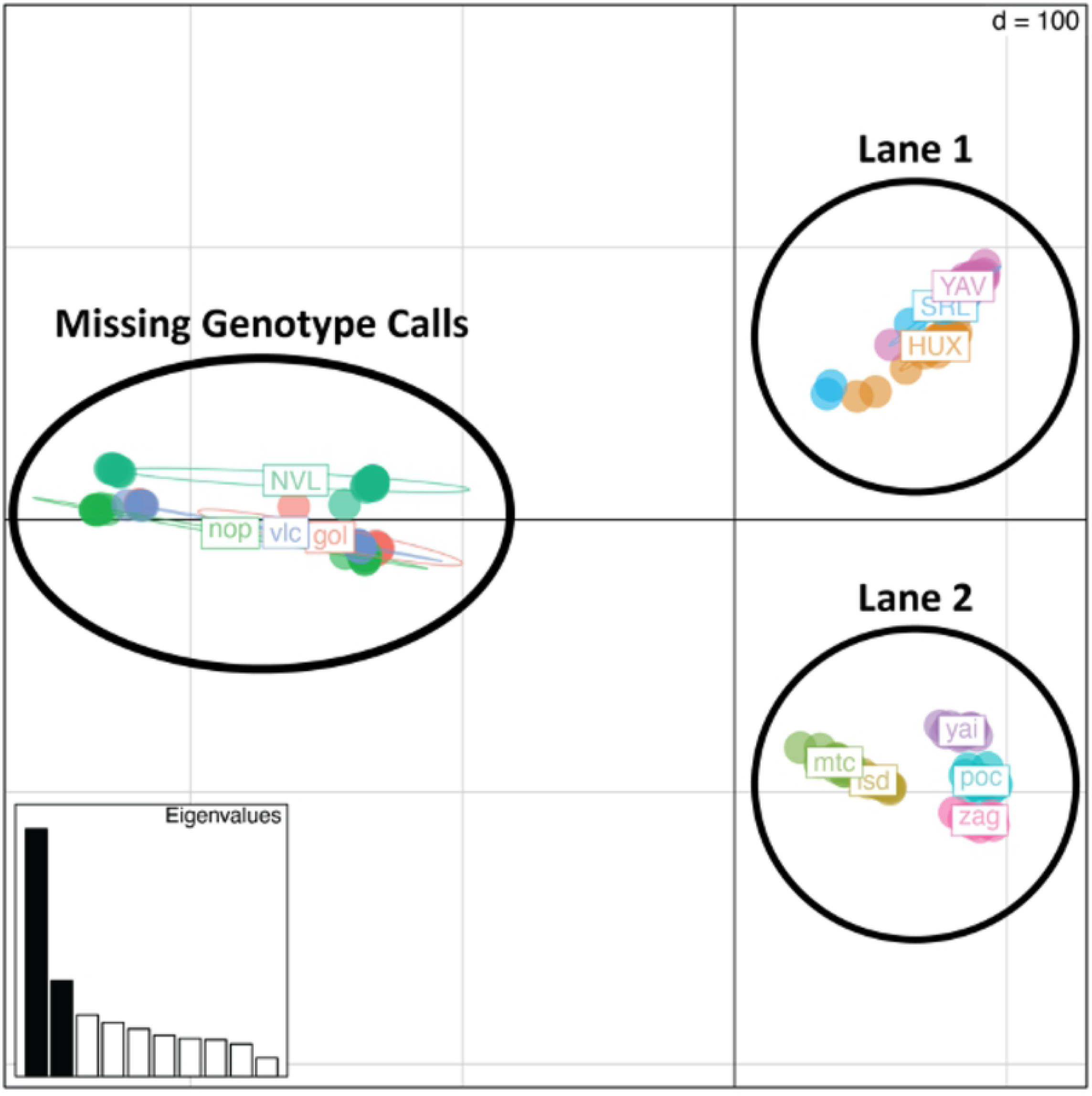
Principal components analysis of genetic variance illustrating the effect of sequencing lane and missing genotype calls for individuals in certain populations. Each population is color coded and labeled. Each effect group is circled and labeled. The four populations missing genotype calls were missing at least 80% of the genotypes output by populations in Stacks.

### Statistical analyses

Plotting principal components 1 and 2 for the filtered SNPs shows that the first principal component explains the difference between the two sequencing lanes (Fig 3). Plotting principal components 2 and 3 removes the lane effect and shows that weevils from each community group tightly together. There is a group of four communities in Oaxaca that share gene flow and cluster together, while the remaining communities are distinct from one another (Fig 4). In addition, principal component 3 can be explained by which state the community is located. The two communities on the bottom of the graph, Huixtla (HUX) and Montecristo (mtc), are in Chiapas, while all other communities are located in Oaxaca (Fig 4). Nei’s pairwise F_ST_ was calculated using the hierfstat package in R (Table 3). All population pairs have negative FST values, which should be interpreted as no difference between populations.

**Table 3.** Nei’s pairwise F_ST_.

|  | HUX | ISD | MTC | POC | SRL | YAI | YAV |
| --- | --- | --- | --- | --- | --- | --- | --- |
| ISD | -0.175 |  |  |  |  |  |  |
| MTC | -0.214 | -0.237 |  |  |  |  |  |
| POC | -0.106 | -0.198 | -0.260 |  |  |  |  |
| SRL | -0.100 | -0.167 | -0.203 | -0.099 |  |  |  |
| YAI | -0.113 | -0.219 | -0.269 | -0.121 | -0.113 |  |  |
| YAV | -0.093 | -0.164 | -0.197 | -0.090 | -0.091 | -0.103 |  |
| ZEG | -0.111 | -0.205 | -0.256 | -0.120 | -0.108 | -0.122 | -0.099 |

Values calculated in the *hierfstat* package in R for the stringently filtered SNP data. Analysis includes 8 populations and 132 individuals.

**Fig 3.**
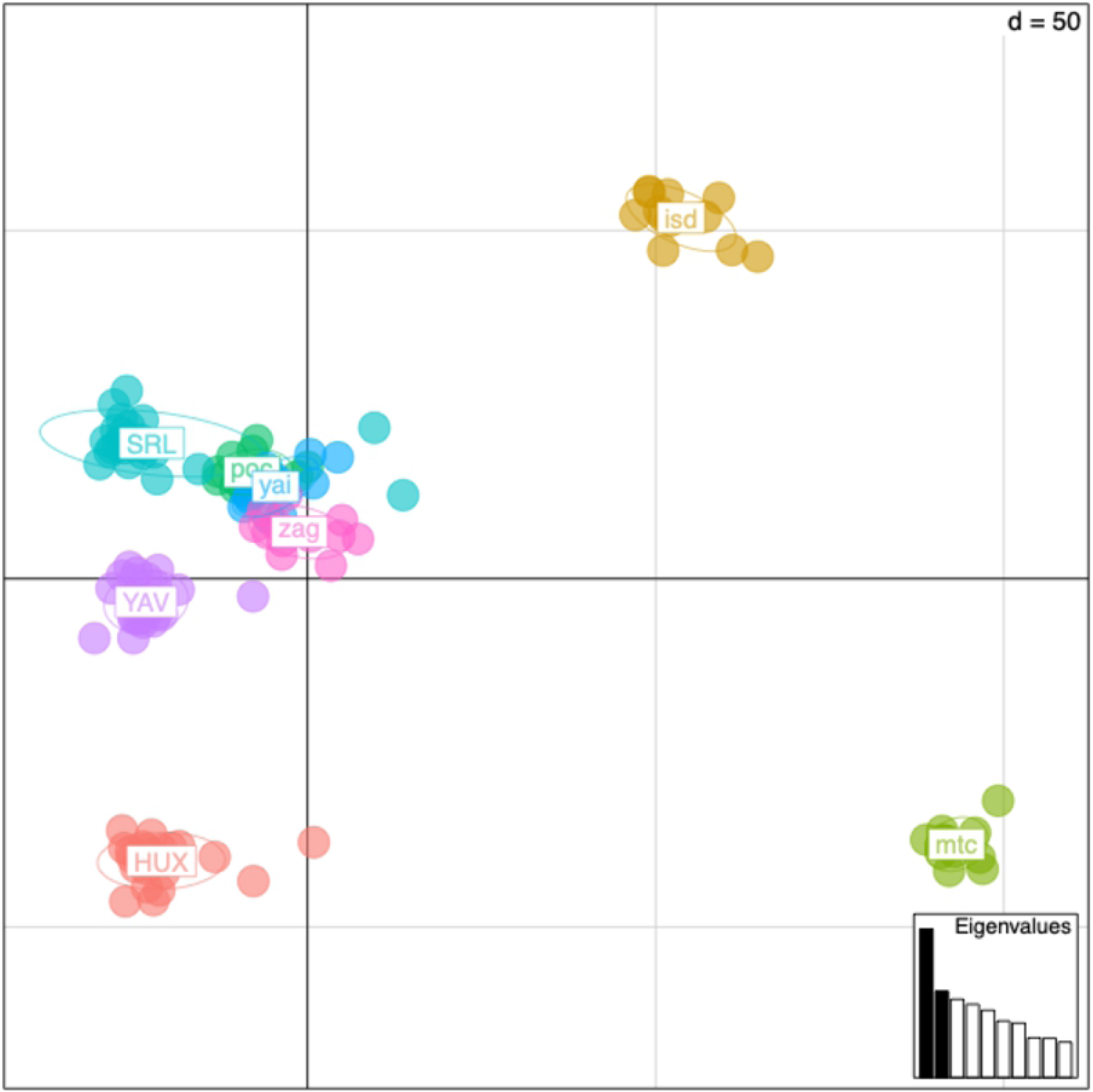
Principal components analysis of genetic variance in filtered data. Weevils from each community are color coded. Principal components 1 and 2 are plotted. Communities SRL, YAV, and HUX are from the first sequencing lane. The remaining populations are from the second sequencing lane. Principal component one is primarily explaining the difference between the two sequencing lanes.

**Fig 4.**
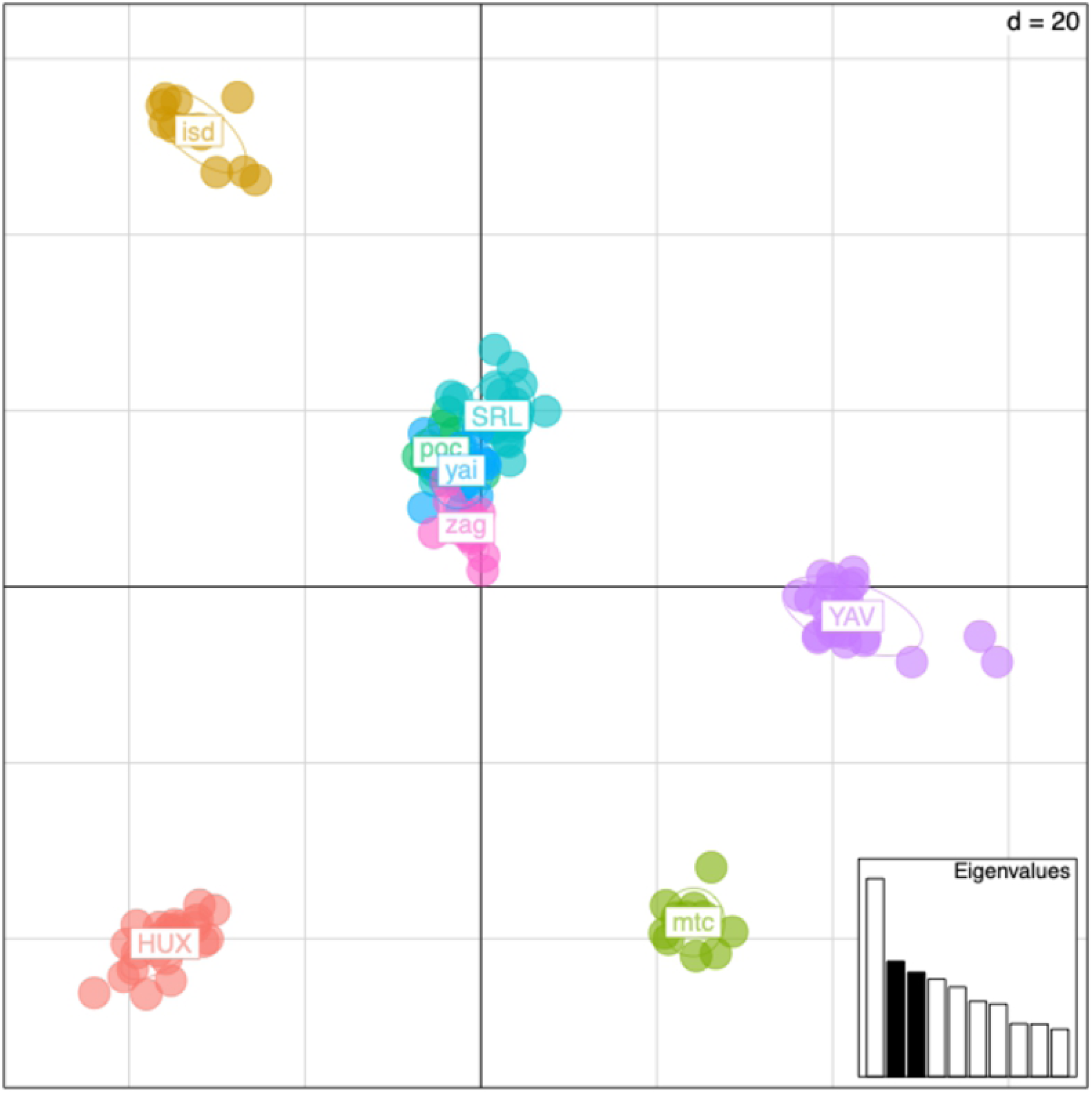
Principal components analysis of genetic variance in filtered data. Weevils from each community are color coded. Principal components 2 and 3 are plotted. Communities SRL, YAV, and HUX are from the first sequencing lane. The remaining populations are from the second sequencing lane.

## Discussion

Maize weevils are a threat to food security in southern and coastal Mexico as they are the primary pest of stored maize. Damage from this pernicious pest typically results in 10% - 40% of postharvest loss, but García-Lara *et al*. [24] reported postharvest losses of up to 100% in southern Mexico, dependent on climate and storage practices. In addition, this region is the center of origin for maize, and subsistence farmers value the landraces that are specifically adapted for their climate and taste preferences [5]. Gaining a better understanding of the existing population genetic structure of this species is important to more responsibly and effectively use traditional and emerging population control techniques aimed at reducing losses from pests.

In this study, we utilized genome-wide SNPs to evaluate the genetic population structure of maize weevil, *S. zeamais*, from the southern and coastal Mexican states of Oaxaca and Chiapas. While other studies have attempted to identify genetic structure of this species at a larger scale, our smaller geographic scale study is the first to find significant genetic population structure in the maize weevil.

Although the negative pairwise F_ST_ results indicate that there is no difference between populations, the principal components analysis was able to identify genetic differentiation based on geography. With the exception of weevils from San Isidro Campechero (ISD) and Santa Maria Yavesia (YAV), the communities in Oaxaca are more genetically more similar to each other than the communities in Chiapas. Principal component 3 explains the difference between communities in Oaxaca and Chiapas. In addition, the communities in Chiapas were more distinct from each other than the 4 clustered communities in Oaxaca. While the exact reasons underlying the genetic structure is unknown, we observed that the roads between certain communities in Oaxaca were easier to traverse and the travel time between sites was subsequently reduced. On the other hand, the communities sampled in Chiapas were much more isolated than those in Oaxaca. Likewise, ISD and YAV, the two genetically distinct communities in Oaxaca were also the most socially isolated communities sampled in that state. The fact that the communities are more connected in Oaxaca than in Chiapas may help to facilitate grain trade between communities, leading to more gene flow between maize weevil populations. An interesting future study could incorporate grain trade routes into the genetic analysis to explore the potential effect of human interactions on the genetic structure of maize weevil. If the effect is found to be significant, trade practices could also inform pest management strategies. An alternative hypothesis to explain the difference between the populations found in Oaxaca and Chiapas is the existence of the Isthmus of Tehuantepec, an extremely windy area of Mexico. This region may form a natural barrier to weevil dispersal if migration is dominated by weevil flight rather than human-mediated grain trade. One or a combination of these factors may be shaping the genetic population structure of maize weevil in southern Mexico.

An important result from this study is the control of sequencing lane effects caused by combining independent sequencing lanes. The effects were controlled by stringent filtering of both individuals and loci. Removing under representative SNPs from the dataset greatly decreased the obvious lane effects. At the same time, a sufficient number of informative SNPs were retained to allow for clear relationships between populations to be observed. While it is preferable to design a ddRadSeq experiment where samples from all populations are randomized across all sequencing lanes, it is reassuring that with proper filtering and conservative SNP selection, this method is robust to identify population structure when randomization cannot be achieved.

This study is the first to characterize significant population structure in the maize weevil. Previous research showed that the maize weevil has experienced a worldwide range expansion over the last several hundred years [7]. Here, we show that although there continues to be gene flow between populations of maize weevil, that fine-scale genetic structure exists. It is possible that this structure is shaped by the movement and trade of maize by humans in the region, by geographic barriers to gene flow, or a combination of factors.

## Acknowledgements

We would like to thank all of the farmers who gave their time, knowledge, and weevils to us. We would also like to thank our guides in Mexico who facilitated our interactions with the farmers. Without their assistance we would not have been able to conduct this study.

## References

1. García-Lara S, Bergvinson DJ. Identification of maize landraces with high level of resistance to storage pests Sitophilus zeamais Motschulsky and Prostephanus truncatus horn in Latin America. Rev Fitotec Mex. 2013;36(Suppl. 3-A): 347–356.

2. Chigoverah AA, Mvumi BM. Efficacy of metal silos and hermetic bags against stored-maize insect pests under simulated smallholder farmer conditions. J Stored Prod Res. 2016;69: 179–189.

3. Oliveira AP, Santos AA, Santana AS, Lima APS, Melo CR, Santana EDR, et al. Essential oil of Lippia sidoides and its major compound thymol: toxicity and walking response of populations of Sitophilus zeamais (Coleoptera: Curculionidae). Crop Prot. 2018;112: 33–38.

4. Arnason JT, Baum B, Gale J, Lambert JDH, Bergvinson D, Philogene BJR, et al. Variation in resistance of Mexican landraces of maize to maize weevil Sitphilus zeamais, in relation to taxonomic and biochemical parameters. Euphytica. 1994;74: 227–236.

5. Hellin J, Bellon MR, Hearne SJ. Maize Landraces and Adaptation to Climate Change in Mexico. J Crop Improv. 2014;28(28): 484–501.

6. Koul O, Walia S, Dhaliwal GS. Essential oils as green pesticides: potential and constraints. Biopestic Int. 2008;4(4): 63–84.

7. Corrêa AS, Vinson CC, Braga LS, Guedes, RNC, de Oliveira LO. Ancient origin and recent range expansion of the maize weevil Sitophilus zeamais, and its genealogical relationship to the rice weevil S. oryzae. Bull Entomol Res. 2017;107(107): 9–20.

8. Ndiaye MR, Sembène M. Genetic structure and phylogeographic evolution of the West African populations of Sitophilus zeamais (Coleoptera, Curculionidae). J Stored Prod Res. 2018;77: 135–143.

9. Giles PH. Observations in Kenya on flight activity of stored products insects particularly Sitophilus zeamais Motsch. J Stored Prod Res. 1969;4(4): 317–329.

10. Morin PA, Martien KK, Taylor BL. Assessing statistical power of SNPs for population structure and conservation studies. Mol Ecol Resour. 2009;9(9): 66–73.

11. Liu N, Chen L, Wang S, Oh C, Zhao H. 2005. Comparison of single-nucleotide polymorphisms and microsatellites in inference of population structure. BMC Genet. 2009;6(Suppl I): S26–5.

12. International Finance Corporation. Investments for a windy harvest: IFC support of the Mexican wind sector drives results. 2014;1: 1–4. Available from: https://www.ifc.org/wps/wcm/connect/a0f55458-988a-4756-8ebd-f456235bc644/IFC_CTF_Mexico.pdf?MOD=AJPERES&CVID=kCCelk9

13. Baltzegar J, Barnes JC, Elsensohn JE, Gutzmann N, Jones MS, King S, et al. Anticipating complexity in the deployment of gene drive insects in agriculture. J Responsible Innov. 2018;5(S1): S81–S97.

14. Peterson BK, Weber JN, Kay EH, Fisher HS, Hoekstra HE. Double Digest RADseq: an inexpensive method for de novo SNP discovery and genotyping in model and non-model species. PLoS One. 2012;7(7): e37135.

15. Fritz ML, Paa S, Baltzegar J, Gould F. Application of a dense genetic map for assessment of genomic responses to selection and inbreeding in Heliothis virescens. Insect Mol Biol. 2016;25(25): 385–400.

16. Andrews S, Krueger F, Segonds-Pichon A, Biggins L, Krueger C, Wingett S, et al. FastQC: a quality control tool for high throughput sequence data. Babraham Bioinformatics. Babraham Institute. 2010-2020. Available from: http://www.bioinformatics.babraham.ac.uk/projects/fastqc.

17. Bolger AM, Lohse M, Usadel B. Trimmomatic: a flexible trimmer for Illumina sequence data. Bioinformatics. 2014;30(30): 2114–2120.

18. Ewels P, Magnusson M, Lundin S, Käller M. MultiQC: summarize analysis results for multiple tools and samples in a single report. Bioinformatics. 2016;32(32): 3047–3048.

19. Rochette NC, Catchen JM. Deriving genotypes from RAD-seq short-read data using Stacks. Nat Protoc. 2017;12: 2640–2659.

20. Catchen J, Hohenlohe PA, Bassham S, Amores A, Cresko WA. Stacks: an analysis tool set for population genomics. Mol Ecol. 2013;22(22): 3124–3140.

21. Jombart T. adegenet: a R package for the multivariate analysis of genetic markers. Bioinformatics. 2008;[accessed 2020 Mar 24];24(11): 1403–1405. Available at: https://academic.oup.com/bioinformatics/article/24/11/1403/191127. doi:10.1093/bioinformatics/btn129.

22. R Project for Statistical Computing. © The R Foundation. 2020. R Foundation for Statistical Computing, Vienna, Austria. Available from: https://www.R-project.org/.

23. Goudet JT. Hierfstat: Estimation and Tests of Hierarchical F-Statistics. 2018. Available from: http://www.r-project.org, http://github.com/jgx65/hierfstat.

24. García-Lara S, García-Jaimes E, Bergvinson DJ. Mapping of maize storage losses due to insect pests in central Mexico. J Stored Prod Res. 2019;84: 101529.

